# Synthesis and In Vitro Characterization of [^3^H]OGA-2506 as a High Affinity Radioligand for O-GlcNAcase

**DOI:** 10.1101/2025.09.19.677343

**Authors:** Yinlong Li, Taoqian Zhao, Zhendong Song, Xin Zhou, Pablo Martinez Pardo, Jiahui Chen, Qilong Hu, Chongjiao Li, Xiaoyan Li, Zhenkun Sun, Yabiao Gao, Danielle E. Hoyle, Jimmy S. Patel, Charles S. Elmore, Hongjie Yuan, Steven H. Liang

**Affiliations:** Department of Radiology and Imaging Sciences, Emory University, 1364 Clifton Road, Atlanta, Georgia 30322, United States; Early Chemical Development, Pharmaceutical Sciences, R&D, AstraZeneca Pharmaceuticals, Gothenburg 43183, Sweden; Department of Pharmacology and Chemical Biology, Emory University School of Medicine, Atlanta, Georgia, 30322, United States; Department of Radiation Oncology, Winship Cancer Institute of Emory University, Atlanta, Georgia, 30322, United States; Early Chemical Development, Pharmaceutical Sciences, R&D, AstraZeneca Pharmaceuticals, Boston, Massachusetts 02451 United States

**Keywords:** O-GlcNAcase, tritium labeling, radioligand, radioligand binding assay, autoradiography

## Abstract

O-GlcNAcase (OGA) is a glycoside hydrolase that regulates protein O-GlcNAcylation, a dynamic post-translational modification implicated in numerous cellular processes. Dysregulation of OGA alters cellular O-GlcNAc homeostasis and has been linked to neurodegenerative and other chronic diseases. The development of radioligands targeting OGA, particularly those derived from well-characterized tool compounds, could substantially advance drug discovery in this area. Herein, we report the radiosynthesis of a novel tritium (^3^H)-labeled radioligand **7** (code name [^3^H]OGA-2506) derived from a chiral piperidine scaffold, and its preliminary *in vitro* binding evaluation. Starting from iodine precursor **8**, palladium-catalyzed tritiation afforded [^3^H]**7** with excellent molar activity (1822 GBq/mmol) and high radiochemical purity (>99%). Saturation binding studies revealed that [^3^H]**7** binds OGA with high affinity to rat striatum homogenates (K_d_ = 3.56 nM; B_max_ = 42.62 nM). Competition assays yielded an IC_50_ of 5.55 nM, and preliminary autoradiography demonstrated heterogeneous regional distribution in the brain with specific binding. These findings highlight [^3^H]**7** as a valuable tool for OGA binding studies and provide a foundation for the future development of novel positron emission tomography (PET) ligands to probe O-GlcNAc signaling in the brain.

## Introduction

O-Linked β-N-acetylglucosaminylation (O-GlcNAcylation) is a critical post-translational modification in which a single *N*-acetylglucosamine (GlcNAc) moiety is added to serine or threonine residues of proteins.^1-3^ This modification is dynamically regulated by O-GlcNAc transferase (OGT), which installs the modification, and O-GlcNAcase (OGA), which removes it.^4, 5^ Through their opposing activities, OGT and OGA maintain the dynamic balance of protein O-GlcNAcylation, thereby modulating key cellular processes such as transcriptional regulation,^6^ protein stability,^7^ signal transduction,^8^ and stress responses.^9^ Dysregulation of O-GlcNAc cycling perturbs cellular homeostasis and has been implicated in various diseases.^10, 11^ For instance, reduced O-GlcNAcylation in Alzheimer’s disease (AD) has been associated with increased tau hyperphosphorylation and neurodegeneration.^12, 13^ Similarly, OGA inhibition has been shown to reduce α-synuclein aggregation and improve motor function in preclinical model of Parkinson’s disease (PD).^14, 15^ Meanwhile, altered OGA activity has been associated with diabetes,^16^ cancer progression,^17^ and cardiovascular disorders.^18^ These findings highlight OGA as a critical therapeutic target across diverse pathological contexts. Over the past decades, numerous OGA inhibitors have been reported.^19^ Early drug discovery efforts identified thiazoline and imidazole scaffolds as promising chemotypes,^20, 21^ with Thiamet G representing a benchmark thiazoline-based OGA inhibitor of high potency.^22^ More recently, piperidine-based scaffolds have been explored, with candidates including MK-8719^23^ and ASN120290^15^ entering clinical evaluation for AD and progressive supranuclear palsy (PSP).^24, 25^ However, no OGA inhibitor has yet been approved, and the development of selective and potent inhibitors with favorable pharmacokinetic properties remains a challenge.

Tritium-labeled radioligands (^3^H-ligands) represent powerful tools for *in vitro* evaluation of enzyme inhibitors, offering high specific activity and precise quantitative detection.^26, 27^ They are extensively used in saturation and competition binding assays and are compatible with autoradiography to map enzyme distribution.^28, 29^ To date, several ^3^H-labeled OGA ligands such as [^3^H]LSN3316612,^30^ [^3^H]CH_3_-BIO-1790735 and [^3^H]BIO-1819578 have been reported (**Figure 1**).^31^ These ligands have greatly facilitated the study of OGA inhibitors and positron emission tomography (PET) tracers. Here, we report the radiosynthesis and preliminary *in vitro* binding characterization of a novel ^3^H-labeled OGA radioligand ([^3^H]**7**) derived from a chiral piperidine scaffold. [^3^H]**7** demonstrates high affinity and selectivity toward OGA, providing an advanced tool for *in vitro* inhibitor characterization and potential PET tracer development.^32, 33^

**Figure 1.**
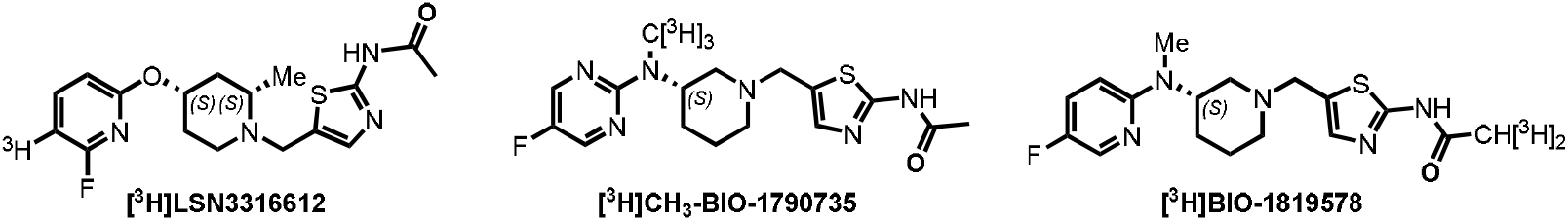
OGA targeted tritium-labeled ligands.

## Materials and methods

### General

Unless otherwise noted, all reagents were obtained from commercial suppliers and used without further purification. Compound **7** (OGA-2506) was prepared following the reported procedure.^34^ Nuclear magnetic resonance (NMR) spectra were recorded on a Bruker AVANCE NEO 400MHZ spectrometer, with chemical shifts (δ, ppm) referenced to tetramethylsilane (TMS) via the residual solvent signal. High-performance liquid chromatography–mass spectrometry (HPLC-MS) was performed on either a Shimadzu LCMS-2010EV quadrupole mass spectrometer with an ESI ion source or an Agilent 6120B quadrupole LC/MS system with an ESI ion source. Analytical HPLC was conducted using a Shimadzu LC-20AB system equipped with a PDA detector set at 220 and 254 nm.

### Chemistry

#### *(S)-((1-(tert-butoxycarbonyl)piperidin-3-yl)methyl)zinc(II) iodide* (2)

To a round-bottom flask was added I_2_ (15.6 mg, 61.5 μmol), DMF (8.00 mL) and Zn (804 mg, 12.3 mmol) at 0 °C under N_2_ atmosphere. Then compound **1** (2.00 g, 6.15 mmol) in DMF (8.00 mL) was added at 0 °C. The resulting mixture was stirred for 2 hrs at 35 °C under N_2_. The reaction mixture was used without further work-up.

#### *tert-butyl (R)-3-(4-methoxybenzyl)piperidine-1-carboxylate* (4)

A mixture of compound **3** (150 mg, 640 μmol), compound **2** (0.23 *M*, 6.97 mL), SPhos (52.6 mg, 128 μmol) and Pd_2_(dba)_3_ (58.7 mg, 64.1 μmol) in DMF (2.00 mL) was degassed and purged with N_2_ for 3 times, stirred at 50 °C for 2 hrs under N_2_ atmosphere. The reaction mixture was partitioned between H_2_O (10.0 mL) and EtOAc (30.0 mL). The organic phase was separated, washed with aqueous NaCl (10.0 mL), dried over Na_2_SO_4_, concentrated under reduced pressure to give a residue. The residue was purified by chromatography on silica gel (0-5% EtOAc/Hexane) to afford compound **4** (153 mg, 450 μmol, 70% yield) as a colorless oil. ^1^H NMR (400 MHz, CDCl_3_) δ 7.07 (d, *J* = 8.4 Hz, 2H), 6.79 - 6.85 (m, 2H), 3.84 - 3.97 (m, 2H), 3.79 (s, 3H), 2.73 - 2.83 (m, 1H), 2.48 -2.56 (m, 2H), 2.36 - 2.43 (m, 1H), 1.60 - 1.79 (m, 3H), 1.44 (s, 9H), 1.34 - 1.41 (m, 1H), 1.05-1.16 (m, 1H). LCMS: m/z = 206.2 (M+H-100)^+^.

#### *(R)-3-(4-methoxybenzyl)piperidine* (5)

A solution of compound **4** (198 mg, 648 μmol) in HCl/dioxane (2 *M*, 3.32 mL) was stirred at 30 °C for 1 hr under N_2_ atmosphere. The reaction mixture was concentrated under reduced pressure to give compound **5** (160 mg) as a yellow solid. ^1^H NMR (400 MHz, DMSO-d_6_) δ 8.29-8.73 (m, 1H), 7.09 (d, *J* = 8.4 Hz, 2H), 6.87 (d, *J* = 8.8 Hz, 2H), 3.72 (s, 3H), 3.17 (br d, *J* = 12.4 Hz, 1H), 3.03 (br d, *J* = 11.6 Hz, 1H), 2.63 - 2.79 (m, 1H), 2.47 (br s, 3H), 1.81 - 1.97 (m, 1H), 1.64 - 1.79 (m, 2H), 1.48 - 1.63 (m, 1H), 1.04 - 1.24 (m, 1H). LCMS: m/z = 206.2 (M+H)^+^.

#### *(R)-N-(5-((3-(4-methoxybenzyl)piperidin-1-yl)methyl)thiazol-2-yl)acetamide* (7)

A mixture of compound **5** (160 mg, 779 μmol), compound **6** (149 mg, 779 μmol), DIEA (302 mg, 2.34 mmol, 407 μL) in DMF (5.00 mL) was degassed and purged with N_2_ for 3 times, stirred at 25 °C for 7 hrs. The mixture was diluted with H_2_O (30.0 mL) and extracted with EtOAc (75.0 mL * 2). The combined organic layers were washed with brine (75.0 mL * 2), dried over Na_2_SO_4_, filtered and concentrated under reduced pressure to give a residue. The residue was purified by prep-HPLC (column: F-Welch Xtimate C18 40*200mm 7 μm;mobile phase: [H_2_O(0.05% NH_3_·H_2_O+10 mM NH_4_HCO_3_)-ACN]; gradient:26%-100% B over 20.0 min) to give compound **7** (147.8 mg, 52% yield) as a white solid. ^1^H NMR (400 MHz, CDCl_3_) δ 11.57 (br s, 1H), 7.17 (s, 1H), 7.04 (d, *J* = 8.8 Hz, 2H), 6.81 (d, *J* = 8.8 Hz, 2H), 3.78 (s, 3H), 3.54 - 3.74 (m, 2H), 2.78 (br d, *J* = 9.2 Hz, 2H), 2.35 - 2.53 (m, 2H), 2.30 (s, 3H), 1.98 (br t, *J* = 10.0 Hz, 1H), 1.74 - 1.87 (m, 2H), 1.61 - 1.67 (m, 2H), 1.46 - 1.54 (m, 1H), 0.83 - 0.99 (m, 1H). LCMS: m/z = 360.3 (M+H)^+^.

#### *(R)-N-(5-((3-(3,5-diiodo-4-methoxybenzyl)piperidin-1-yl)methyl)thiazol-2-yl)acetamide* (8)

A solution of compound **7** (30.0 mg, 83.5 μmol), NIS (56.3 mg, 250 μmol) in TFA (3.00 mL) was stirred at 25 °C for 20 min under N_2_. The mixture was diluted with H_2_O (5.00 mL) and NaHCO_3_ (20.0 mL). The mixture was extracted with CH_2_Cl_2_ (25.0 mL * 2). The combined organic layers were washed with brine (30.0 mL * 2), dried over Na_2_SO_4_, filtered and concentrated under reduced pressure to give a residue. The residue was purified by prep-HPLC (column: F-Welch Xtimate C18 40*200mm 7 μm; mobile phase: [H_2_O (0.225% FA)-ACN]; gradient: 2%-42% B over 20.0 min) to give compound **8** (40.0 mg, 77 % yield) as a white solid. ^1^H NMR (400 MHz, CDCl_3_) δ 10.51-11.54 (m, 1H), 7.52 (s, 2H), 7.21 (br s, 1H), 3.82 (s, 3H), 3.68 (br d, *J* = 7.0 Hz, 2H), 2.74 - 2.84 (m, 2H), 2.40 - 2.47 (m, 1H), 2.30 (s, 3H), 2.01 - 2.10 (m, 2H), 1.74-1.83 (m, 2H), 1.66 (br d, *J* = 7.6 Hz, 2H), 1.57 (br d, *J* = 2.8 Hz, 1H), 0.93 (br d, *J* = 10.0 Hz, 1H). LCMS: m/z = 611.9 (M+H)^+^.

### Radiochemistry

In a reaction flask, palladium on carbon 10% (1.7 mg, 1.64 µmol) was loaded and a solution of compound **8** (1 mg, 1.64 µmol) in MeOH (0.5 mL) was added. The system was cooled down to - 196 °C and it was degassed three times. Tritium (1040 mbar) was loaded, and the reaction was allowed to warm up to room temperature and stir for 3 hours. The catalyst was removed by filtration, and the solvent was evaporated. The residue was purified by HPLC (Waters Xbridge C18, 5 µm column). The product containing fractions was combined, and the solvent was evaporated. The residue was dissolved in 5 mL of ethanol to give 750 MBq of [^3^H]**7**. The radiochemical purity was determined to be > 99.5% with a UV area % (254 nm) of 100%. The molar activity was determined to be 1822 GBq/mmol. LC/MS: 360.1 (3.0%), 362.2 (31.9%), 364.2 (100%), 365.3 (17.8%), 366.3 (7.6%).

### Radioligand binding assay

Frozen tissue was homogenized in lysis buffer (50 mM Tris-HCl, 5 mM MgCl_2_, 5 mM EDTA, protease inhibitors). After low-speed centrifugation (100 × g, 3 min) to remove debris, membranes were pelleted by centrifugation (17,000 × g, 10 min, 4 °C), washed once, and resuspended in buffer with 10% sucrose. Aliquots were stored at –80 °C, and protein concentration was determined using the Pierce® BCA assay. Binding assays were performed in 96-well plates (250 µL final volume) with membranes (3–20 µg for cells; 50–120 µg for tissue), radioligand (0.2–20 nM), and buffer or unlabeled ligand (to define non-specific binding). Incubations were carried out at 30 °C for 60 min and terminated by filtration through 0.3% PEI-soaked GF/C filters. Filters were washed with cold buffer, dried, and counted in a scintillation counter. Specific binding was calculated as total minus non-specific binding. Data were fitted by non-linear regression (saturation binding) in GraphPad Prism to obtain K_d_ and B_max_ values.

### *In vitro* autoradiography

Briefly, frozen tissue sections were thawed at room temperature for 15 min and pre-rinsed sequentially in 50 mM Tris-HCl (pH 7.4, 15 min) and incubation buffer (15 min; saline containing 0.2% BSA, 0.01% Tween-20, 1% ethanol, 1% DMSO). For nonspecific binding, 10 µM thiamet-G was included. Sections were then incubated for 2 h at room temperature in incubation buffer containing 25 nM of [^3^H]**7**, followed by washes in incubation buffer, saline with 0.2% BSA, saline, and ice-cold water (2 min each). Slides were dried (30 min) and exposed to paraformaldehyde vapor under vacuum (10 min), and scanned using a BeaQuant-S system.

## RESULTS AND DISCUSSION

### Chemistry

As depicted in **Scheme 1**, the target compound **7** was synthesized from the commercially available tert-butyl (*S*)-3-(iodomethyl)piperidine-1-carboxylate (**1**). Reduction of compound **1** with zinc and iodine afforded the corresponding organozinc intermediate **2**, which was subsequently coupled with aryl iodide **3** *via* Negishi cross-coupling reaction to give compound **4** in 70% yield. Deprotection of the Boc group using HCl produced the free amine **5**, which was then reacted with chloride **6** to furnish the desired compound **7** in 52% yield over two steps.

**Scheme 1.**
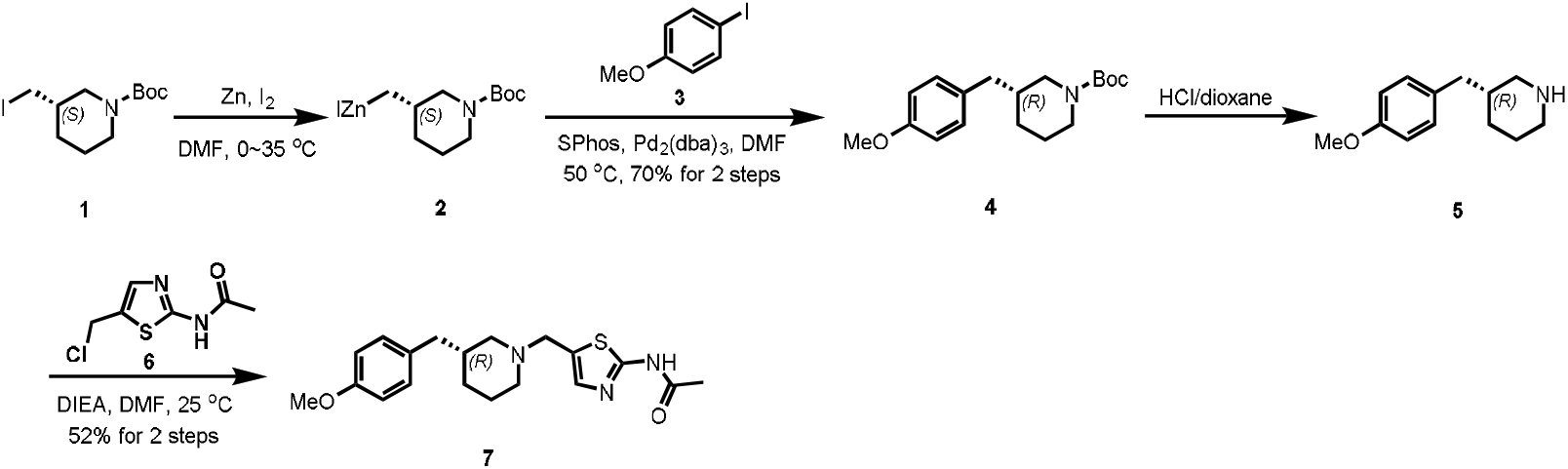
Synthesis of compound **7**.

### Physicochemical and pharmacological properties

The *in silico* physicochemical properties of compound **7** were predicted using ChemDraw and ACD/Labs, and the results are summarized in Table 1. Compound **7** has a molecular weight (MW) of 359.49, a calculated logP (clogP) of 3.28, and a topological polar surface area (TPSA) of 53.93 Å^2^, all of which are consistent with favorable central nervous system (CNS) penetration. The presence of a single hydrogen bond donor (HBD = 1) further supports brain permeability in accordance with Lipinski’s rule of five. In addition, the predicted brain-to-blood partition coefficient (LogBB) of 0.06 and a CNS multiparameter optimization (MPO) score of 5.2 suggest good CNS drug-like properties, thereby supporting the potential of compound **7** for further evaluation.

**Table 1.** ^*a*^Values were calculated with ChemDraw 21.0 software. ^*b*^Values were predicted with ACD/labs.

| MW<sup>a</sup> | clogP<sup>a</sup> | tPSA<sup>a</sup> | HBD<sup>b</sup> | Lipinski's rule<sup>b</sup> | LogBB<sup>b</sup> | MPO score<sup>b</sup> |
| --- | --- | --- | --- | --- | --- | --- |
| 359.49 | 3.28 | 53.93 | 1 | 0 | 0.06 | 5.2 |

### Radiochemistry

The iodine precursor **8** was designed to enable the synthesis of [^3^H]**7** through Pd-catalyzed tritiation. In brief, selective diiodination of compound **7** with *N*-iodosuccinimide (NIS) in trifluoroacetic acid (TFA) furnished the diiodinated intermediate **8** in 77% yield (**Scheme 2**). Subsequent catalytic tritiation of **8** with tritium gas (T_2_) over Pd/C under mild conditions afforded the tritiated radioligand [^3^H]**7**. The resulting [^3^H]**7** was obtained with high radiochemical purity (>99.5%) and high molar activity (1822 GBq/mmol), confirming its suitability for *in vitro* binding studies and autoradiography experiments.

**Scheme 2.**
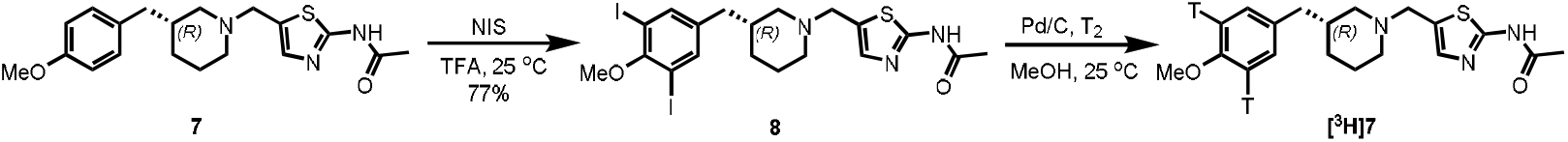
Synthesis of precursor **8** and subsequent radiosynthesis of [^3^H]**7**.

### *In vitro* binding profile

Subsequent *in vitro* binding studies of [^3^H]**7** in rat striatum homogenates demonstrated high-affinity binding (K_d_ = 3.56 nM, B_max_ = 42.62 nM). Competition assays determined an IC_50_ of 5.55 nM, and the calculated binding potential (B_max_/K_d_ > 10) indicates strong specific binding (**Figure 2**). These results establish [^3^H]**7** as a robust radioligand suitable for autoradiography and inhibitor screening applications.

**Figure 2.**
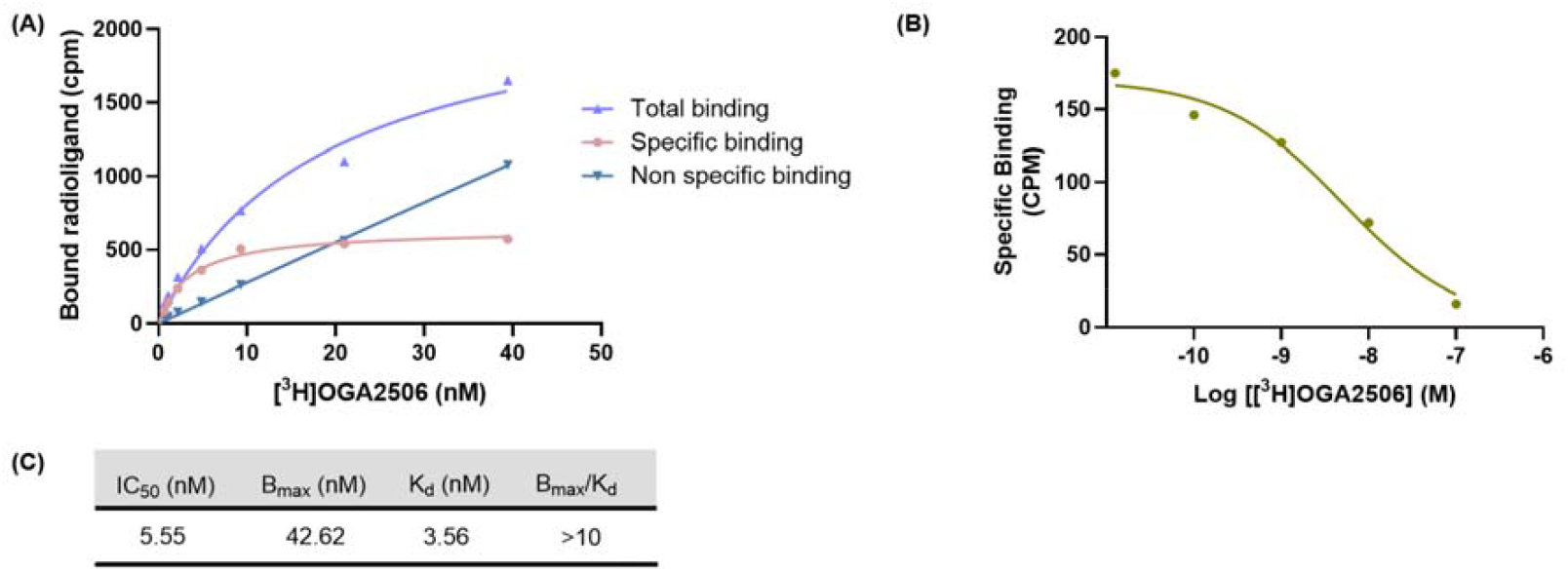
Characterization of [^3^H]**7** binding to rat striatum homogenate. (A) Saturation binding curve of [^3^H]**7** at different concentrations. (B) Competitive binding assay curve of [^3^H]**7** to determine its binding affinity. (C) Summarized binding parameters of [^3^H]**7**.

### *In vitro* autoradiography

*In vitro* autoradiography was performed in rat brain sections to assess the binding selectivity and regional distribution of [^3^H]**7**. As shown in **Figure 3**, [^3^H]**7** displayed heterogeneous distribution, with high binding in the striatum, cortex, and hippocampus, and minimal uptake in the cerebellum. This distribution pattern is consistent with previously reported OGA expression profiles. Co-incubation with 10 μM Thiamet-G, a validated OGA-specific inhibitor, markedly reduced radioligand signal across all regions, confirming specific binding. These findings highlight [^3^H]**7** as a promising ligand for the development of OGA-targeted PET tracers.

**Figure 3.**
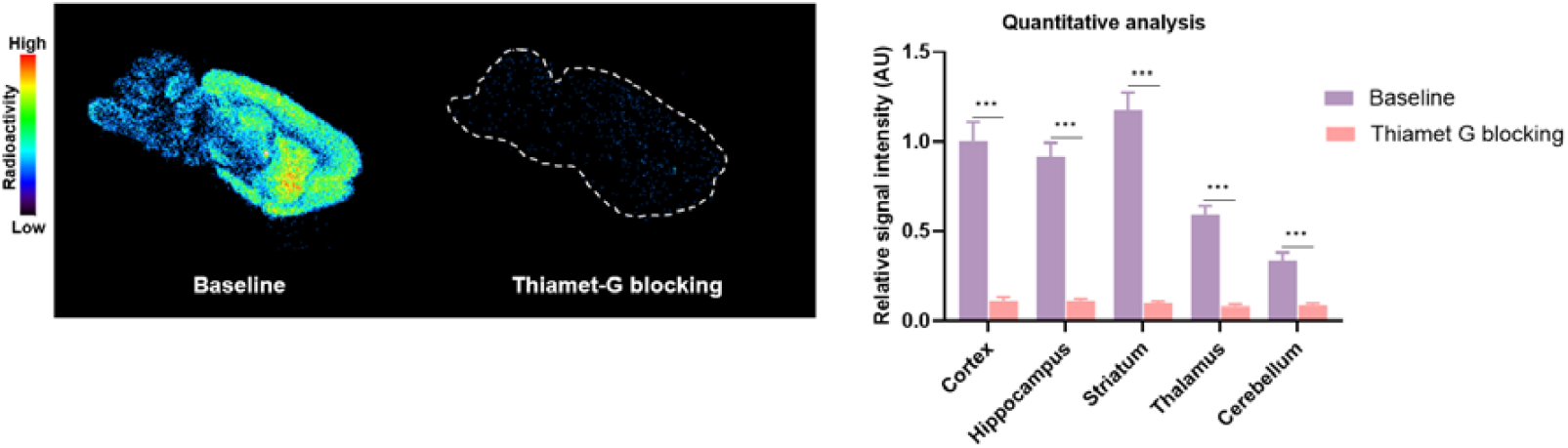
Representative *in vitro* autoradiograms and quantification of [^3^H]**7** in SD rat sagittal brain sections under baseline and blocking conditions. \*\*\**p* ≤ 0.001. Data are shown as mean ± SD.

## CONCLUSION

In this study, we developed a novel OGA-targeted ligand [^3^H]**7** and systematically evaluated its *in vitro* binding properties. Saturation and competition binding assays in rat striatum homogenates demonstrated high-affinity binding and specific interaction with OGA. *In vitro* autoradiography of rat brain sections exhibited heterogeneous distribution, with high uptake in regions such as the striatum, cortex, and hippocampus, consistent with reported OGA expression and confirmed target specificity. Collectively, these *in vitro* results underscore the utility of [^3^H]**7** in OGA research, with potential contributions to future OGA-targeted PET tracer development.

## ACKNOWLEDGMENTS

We thank the Department of Radiology and Imaging Sciences at Emory University School of Medicine for general support. We also acknowledge the technical assistance of Rémy Sence and Jérôme Donnard from Ai4r for their support with the BeaQuant-S system.

## Reference

1. Yang, X.; Qian, K., Protein O-GlcNAcylation: emerging mechanisms and functions. Nature Reviews Molecular Cell Biology 2017, 18 (7), 452–465.

2. Ma, J.; Wu, C.; Hart, G. W., Analytical and Biochemical Perspectives of Protein O-GlcNAcylation. Chemical Reviews 2021, 121 (3), 1513–1581.

3. Hurtado-Guerrero, R.; Dorfmueller, H. C.; van Aalten, D. M. F., Molecular mechanisms of O-GlcNAcylation. Current Opinion in Structural Biology 2008, 18 (5), 551–557.

4. Ma, J.; Hart, G. W., O-GlcNAc profiling: from proteins to proteomes. Clinical Proteomics 2014, 11 (1), 8.

5. Nagel, A. K.; Ball, L. E., O-GlcNAc transferase and O-GlcNAcase: achieving target substrate specificity. Amino Acids 2014, 46 (10), 2305–2316.

6. Parker, M. P.; Peterson, K. R.; Slawson, C., O-GlcNAcylation and O-GlcNAc Cycling Regulate Gene Transcription: Emerging Roles in Cancer. 2021, 13 (7), 1666.

7. Chu, C.-S.; Lo, P.-W.; Yeh, Y.-H.; Hsu, P.-H.; Peng, S.-H.; Teng, Y.-C.; Kang, M.-L.; Wong, C.-H.; Juan, L.-J., O-GlcNAcylation regulates EZH2 protein stability and function. Proceedings of the National Academy of Sciences 2014, 111 (4), 1355–1360.

8. Ong, Q.; Han, W.; Yang, X., O-GlcNAc as an Integrator of Signaling Pathways. Frontiers in Endocrinology 2018, 9, 599.

9. Liu, Y.; Yao, R.-Z.; Lian, S.; Liu, P.; Hu, Y.-J.; Shi, H.-Z.; Lv, H.-M.; Yang, Y.-Y.; Xu, B.; Li, S.-Z., O-GlcNAcylation: the “stress and nutrition receptor” in cell stress response. Cell Stress and Chaperones 2021, 26 (2), 297–309.

10. Wenzel, D. M.; Olivier-Van Stichelen, S., The O-GlcNAc cycling in neurodevelopment and associated diseases. Biochemical Society Transactions 2022, 50 (6), 1693–1702.

11. Bond, M. R.; Hanover, J. A., O-GlcNAc Cycling: A Link Between Metabolism and Chronic Disease. 2013, 33, 205–229.

12. Liu, F.; Shi, J.; Tanimukai, H.; Gu, J.; Gu, J.; Grundke-Iqbal, I.; Iqbal, K.; Gong, C.-X., Reduced O-GlcNAcylation links lower brain glucose metabolism and tau pathology in Alzheimer’s disease. Brain 2009, 132 (7), 1820–1832.

13. Sharma, A.; Singh, A.; Debnath, R.; Gupta, G. D.; Sharma, K., Role of O-GlcNAcylation in Alzheimer’s disease: Insights and perspectives. European Journal of Medicinal Chemistry Reports 2024, 12, 100195.

14. Balana, A. T.; Mahul-Mellier, A.-L.; Nguyen, B. A.; Horvath, M.; Javed, A.; Hard, E. R.; Jasiqi, Y.; Singh, P.; Afrin, S.; Pedretti, R.; Singh, V.; Lee, V. M. Y.; Luk, K. C.; Saelices, L.; Lashuel, H. A.; Pratt, M. R., O-GlcNAc forces an α-synuclein amyloid strain with notably diminished seeding and pathology. Nature Chemical Biology 2024, 20 (5), 646–655.

15. Permanne, B.; Sand, A.; Ousson, S.; Nény, M.; Hantson, J.; Schubert, R.; Wiessner, C.; Quattropani, A.; Beher, D., O-GlcNAcase Inhibitor ASN90 is a Multimodal Drug Candidate for Tau and α-Synuclein Proteinopathies. ACS Chemical Neuroscience 2022, 13 (8), 1296–1314.

16. Forsythe, M. E.; Love, D. C.; Lazarus, B. D.; Kim, E. J.; Prinz, W. A.; Ashwell, G.; Krause, M. W.; Hanover, J. A., Caenorhabditis elegans ortholog of a diabetes susceptibility locus: oga-1 (O-GlcNAcase) knockout impacts O-GlcNAc cycling, metabolism, and dauer. Proceedings of the National Academy of Sciences 2006, 103 (32), 11952–11957.

17. Hu, C.-W.; Wang, A.; Fan, D.; Worth, M.; Chen, Z.; Huang, J.; Xie, J.; Macdonald, J.; Li, L.; Jiang, J., OGA mutant aberrantly hydrolyzes O-GlcNAc modification from PDLIM7 to modulate p53 and cytoskeleton in promoting cancer cell malignancy. Proceedings of the National Academy of Sciences 2024, 121 (24), e2320867121.

18. Masaki, N.; Feng, B.; Bretón□Romero, R.; Inagaki, E.; Weisbrod, R. M.; Fetterman, J. L.; Hamburg, N. M., O□GlcNAcylation Mediates Glucose□Induced Alterations in Endothelial Cell Phenotype in Human Diabetes Mellitus. Journal of the American Heart Association 2020, 9 (12), e014046.

19. Bartolomé-Nebreda, J. M.; Trabanco, A. A.; Velter, A. I.; Buijnsters, P., O-GlcNAcase inhibitors as potential therapeutics for the treatment of Alzheimer’s disease and related tauopathies: analysis of the patent literature. Expert Opinion on Therapeutic Patents 2021, 31 (12), 1117–1154.

20. Cheng, S. S.; Mody, A. C.; Woo, C. M., Opportunities for Therapeutic Modulation of O-GlcNAc. Chemical Reviews 2024, 124 (22), 12918–13019.

21. Sun, S.; Cao, J.; Ji, S.; Wang, J., Recent Progress of Small-molecule Inhibitors of O-GlcNAcase for Alzheimer’s Disease. Mini-Reviews in Medicinal Chemistry 2025. Doi: 10.2174/0113895575376839250606183944

22. Yuzwa, S. A.; Macauley, M. S.; Heinonen, J. E.; Shan, X.; Dennis, R. J.; He, Y.; Whitworth, G. E.; Stubbs, K. A.; McEachern, E. J.; Davies, G. J.; Vocadlo, D. J., A potent mechanism-inspired O-GlcNAcase inhibitor that blocks phosphorylation of tau in vivo. Nature Chemical Biology 2008, 4 (8), 483–490.

23. Selnick, H. G.; Hess, J. F.; Tang, C.; Liu, K.; Schachter, J. B.; Ballard, J. E.; Marcus, J.; Klein, D. J.; Wang, X.; Pearson, M.; Savage, M. J.; Kaul, R.; Li, T.-S.; Vocadlo, D. J.; Zhou, Y.; Zhu, Y.; Mu, C.; Wang, Y.; Wei, Z.; Bai, C.; Duffy, J. L.; McEachern, E. J., Discovery of MK-8719, a Potent O-GlcNAcase Inhibitor as a Potential Treatment for Tauopathies. Journal of Medicinal Chemistry 2019, 62 (22), 10062–10097.

24. Smith, S. M.; Struyk, A.; Jonathan, D.; Declercq, R.; Marcus, J.; Toolan, D.; Wang, X.; Schachter, J. B.; Cosden, M.; Pearson, M.; Hess, F.; Selnick, H.; Salinas, C.; Li, W.; Duffy, J.; McEachern, E.; Vocadlo, D.; Renger, J. J.; Eric, H. D.; Forman, M.; Schoepp, D., O2-13-04: Early Clinical Results and Preclinical Validation of the O-Glcnacase (OGA) Inhibitor Mk-8719 as a Novel Therapeutic for the Treatment of Tauopathies. Alzheimer’s & Dementia 2016, 12 (7S_Part_5), P261–P261.

25. Ryan, J. M.; Quattropani, A.; Abd-Elaziz, K.; den Daas, I.; Schneider, M.; Ousson, S.; Neny, M.; Sand, A.; Hantson, J.; Permanne, B.; Wiessner, C.; Beher, D., O1-12-05: Phase 1 study in healthy volunteers of the O-GlcNAcase inhibitor ASN120290 as a novel therapy for progressive supranuclear palsy and related tauopathies. Alzheimer’s & Dementia 2018, 14 (7S_Part_4), P251–P251.

26. Teng, Y.; Yang, H.; Tian, Y., The Development and Application of Tritium-Labeled Compounds in Biomedical Research. 2024, 29 (17), 4109.

27. Atzrodt, J.; Derdau, V.; Kerr, W. J.; Reid, M., Deuterium- and Tritium-Labelled Compounds: Applications in the Life Sciences. Angewandte Chemie International Edition 2018, 57 (7), 1758–1784.

28. Lever, S. Z.; Fan, K.-H.; Lever, J. R., Tactics for preclinical validation of receptor-binding radiotracers. Nuclear Medicine and Biology 2017, 44, 4–30.

29. Maguire, J. J.; Kuc, R. E.; Davenport, A. P., Radioligand Binding Assays and Their Analysis. In Receptor Binding Techniques, Davenport, A. P., Ed. Humana Press: Totowa, NJ, 2012; pp 31–77.

30. Lu, S.; Haskali, M. B.; Ruley, K. M.; Dreyfus, N. J. F.; DuBois, S. L.; Paul, S.; Liow, J.-S.; Morse, C. L.; Kowalski, A.; Gladding, R. L.; Gilmore, J.; Mogg, A. J.; Morin, S. M.; Lindsay-Scott, P. J.; Ruble, J. C.; Kant, N. A.; Shcherbinin, S.; Barth, V. N.; Johnson, M. P.; Cuadrado, M.; Jambrina, E.; Mannes, A. J.; Nuthall, H. N.; Zoghbi, S. S.; Jesudason, C. D.; Innis, R. B.; Pike, V. W., PET ligands [^18^F]LSN3316612 and [^11^C]LSN3316612 quantify O-linked-β-N-acetyl-glucosamine hydrolase in the brain. Science Translational Medicine 2020, 12 (543), eaau2939.

31. Cook, B. E.; Nag, S.; Arakawa, R.; Lin, E. Y.-S.; Stratman, N.; Guckian, K.; Hering, H.; Lulla, M.; Choi, J.; Salinas, C.; Genung, N. E.; Morén, A. F.; Bolin, M.; Boscutti, G.; Plisson, C.; Martarello, L.; Halldin, C.; Kaliszczak, M. A., Development of a PET Tracer for OGA with Improved Kinetics in the Living Brain. Journal of Nuclear Medicine 2023, 64 (10), 1588.

32. Rong, J.; Haider, A.; Jeppesen, T. E.; Josephson, L.; Liang, S. H., Radiochemistry for positron emission tomography. Nature Communications 2023, 14 (1), 3257.

33. Deng, X.; Rong, J.; Wang, L.; Vasdev, N.; Zhang, L.; Josephson, L.; Liang, S. H., Chemistry for Positron Emission Tomography: Recent Advances in ^11^C-, ^18^F-, ^13^N-, and ^15^O-Labeling Reactions. Angewandte Chemie International Edition 2019, 58 (9), 2580–2605.

34. Genung, N.; Guckian, K. M.; Vessels, J.; Zhang, L.; Gianatassio, R.; Lin, E. Y. S.; Xin, Z. O-glycoprotein-2-acetamido-2-deoxy-3-d-glycopyranosidase inhibitors. WO2019178191A1.

